# Engineering *in vitro* models of cystic fibrosis lung disease using neutrophil extracellular trap inspired biomaterials

**DOI:** 10.1101/2023.06.26.546583

**Authors:** Allison M. Boboltz, Sydney Yang, Gregg A. Duncan

## Abstract

Cystic fibrosis (CF) is a muco-obstructive lung disease where inflammatory responses due to chronic infection result in the accumulation of neutrophil extracellular traps (NETs) in the airways. NETs are web-like complexes comprised mainly of decondensed chromatin that function to capture and kill bacteria. Prior studies have established excess release of NETs in CF airways increases viscoelasticity of mucus secretions and reduces mucociliary clearance. Despite the pivotal role of NETs in CF disease pathogenesis, current *in vitro* models of this disease do not account for their contribution. Motivated by this, we developed a new approach to study the pathobiological effects of NETs in CF by combining synthetic NET-like biomaterials, composed of DNA and histones, with an *in vitro* human airway epithelial cell culture model. To determine the impact of synthetic NETs on airway clearance function, we incorporated synthetic NETs into mucin hydrogels and cell culture derived airway mucus to assess their rheological and transport properties. We found that the addition of synthetic NETs significantly increases mucin hydrogel and native mucus viscoelasticity. As a result, mucociliary transport *in vitro* was significantly reduced with the addition of mucus containing synthetic NETs. Given the prevalence of bacterial infection in the CF lung, we also evaluated the growth of *Pseudomonas aeruginosa* in mucus with or without synthetic NETs. We found mucus containing synthetic NETs promoted microcolony growth and prolonged bacterial survival. Together, this work establishes a new biomaterial enabled approach to study innate immunity mediated airway dysfunction in CF.

## Introduction

The mucus gel layer is one of the first lines of defense to protect the airways by capturing inhaled particles, viruses, and bacteria.^1^ Mucus is continually cleared away by the coordinated beating of cilia on the surface of the airway epithelium, called mucociliary transport (MCT).^2^ However in muco-obstructive lung diseases such as cystic fibrosis (CF), MCT is impaired by alterations to mucus composition that increase the viscoelasticity of the gel.^3,4^ CF is a chronic disease primarily affecting the airways caused by genetic mutation of the cystic fibrosis transmembrane conductance regulator (CFTR).^5^ Neutrophilic lung inflammation is characteristic of CF, and one of the most distinctive changes to airway mucus composition in CF is the over-production of neutrophil extracellular traps (NETs).^5,6^ Neutrophils attack pathogens through a variety of mechanisms, including via the release of NETs in an innate immune response called NETosis.^7^ NETs are webs-like complexes consisting of a porous scaffold of decondensed chromatin with antimicrobial proteins that can expand 3 to 5 times the size of the condensed chromatin.^8^ NETs function to trap and kill bacteria to prevent the spread of infection.^9^ In CF, neutrophils are recruited to the lung in response to rampant bacterial growth in the airways, and undergo NETosis to contain the microbes.^10,11^ However, profuse NETosis has been implicated in several aspects of CF pathophysiology, including increasing mucus viscoelasticity and impairing MCT.^6,12,13^ Counterintuitively, NETs released in excess can also promote bacterial survival and infection in species such as *Pseudomonas aeruginosa*.^12^ *P. aeruginosa* are the most common bacteria infecting CF airways, with a prevalence up to 80% in adult CF patients.^14^ Evidence suggests that *P. aeruginosa* uses the DNA-rich network of NETs to form biofilms and enhance survival.^15,16^

The current gold standard pre-clinical CF model are human airway epithelial (HAE) cell cultures.^17,18^ Primary CF HAE cells can be harvested from explanted tissue and differentiated at an air-liquid interface into a pseudostratified epithelium with characteristics similar to the airway *in vivo*. HAE cell culture models were pivotal in the development of CFTR modulator drugs, which can slow declining lung function and dramatically improve CF patient quality of life.^18–20^ Despite this progress, CF remains an incurable disease and lung function still declines over time even with CFTR modulator treatment, partially due to chronic bacterial infections and persistent inflammation.^20^ Inhaled therapeutics, including antimicrobial agents such as tobramycin and mucolytics such as DNase are commonly used in CF to treat infections and resolve airway obstruction, respectively.^5^ Potentially curative inhaled gene therapies are also being explored as targeted CF therapies.^17,21^ Both *ex vivo* and *in vitro* models have been essential in testing new therapeutics and studying host-pathogen interactions in the CF airway microenvironment.^18,22–24^ Despite our growing understanding of the importance of NET-mediated dysfunction in CF, traditional human *in vitro* models do not account for their presence.

Past work has aimed to address the contributions of NETs in CF pathobiology and their direct impact on the functional properties of mucus. For example, a previous study by Linssen et al examined changes in the macro- and micro-rheological properties of healthy human sputum samples *ex vivo* after incorporation of NETs and found modest changes in the sputum viscoelasticity and microstructure.^25^ However, the study was limited to using small volumes of human sputum samples and the concentrations of NETs harvested through stimulation of neutrophils proved challenging to control. In addition, the process of separating NETs from neutrophils after stimulation can cause protein loss that could compromise the structure of the NETs.^26^ Though sputum samples expectorated by individuals with CF contain NETs, NET and extracellular DNA concentration can vary substantially from patient to patient.^27,28^ HAE cells derived from CF donors produce minimal extracellular DNA and are unable to produce NETs without co-culture and stimulation of neutrophils.^18,29^

To overcome these limitations, we developed a new approach to model the alterations to mucus in CF and recapitulate the impacts of NETs on disease pathogenesis. To do this, we used a synthetic NET (sNET) biomaterial based on previous work to mimic the chromatin scaffold of endogenous NETs.^26,30,31^ This cell-free protocol can be used to generate NET-like biomaterials without the difficulties associated with *in vitro* generation and isolation of endogenous NETs or co-culture of neutrophils within the HAE model. To understand the impact of NETs on the viscoelastic properties of mucus, we incorporated the sNETs into a synthetic mucus hydrogel as described in our prior work^32–34^ and mucus produced by HAE cells. Studies on the functional impact of sNETs on MCT and bacterial infection were also conducted. Our results indicate sNETs can be used to model NET-mediated disease mechanisms *in vitro* and in the future, could be used to explore new therapeutic approaches for CF.

## Experimental

### Synthetic Neutrophil Extracellular Trap (sNET) Preparation

Synthesis of sNETs is based on a previously published protocol, with some minor adaptations.^26,30,31^ DNA (Sigma-Aldrich; sodium salt from salmon testes) was stirred for 2 hours at a concentration of 6 mg/ml in a physiological buffer consisting of 154 mM NaCl, 3 mM CaCl_2_, and 15 mM NaH_2_PO_4_ at pH 7.4. Histones from calf thymus (Sigma-Aldrich; Type II-A) were dissolved separately in the same buffer at a concentration of 6 mg/ml. Equal volumes of the 6 mg/ml DNA and histone solutions were combined where they initially form into a large macroscopic complex. To form a suspension of sNETs at a concentration of 3 mg/ml, ultrasonication was used (Qsonica Q125) for 30 seconds at 20% amplitude. To confirm the formation of sNETs, the suspension was centrifuged for 10 minutes at 15,000 g to form a pellet, then the supernatant was collected. The DNA concentration and A260/280 purity ratio of the supernatant was measured using a spectrophotometer (Nanodrop 2000C, Thermo Scientific). The percent conversion to sNETs was calculated using the following formula: 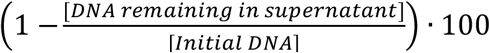. Charge of the sNETs was measured by creating a suspension at a concentration of 50 μg/ml, then using phase analysis light scattering (PALS) to determine the zeta potential with a Nanobrook Omni Particle Analyzer (Brookhaven Instruments).

### Fluorescence Microscopy of sNETs

To image the DNA scaffold structure of sNETs, 20 μl of a suspension of sNETs prepared at a concentration of 50 μg/ml was placed in a microscopy chamber created with an O-ring adhered to a glass microscope slide using vacuum grease. The sNET suspension dried overnight onto the slide, then was stained with 0.01% DAPI solution and washed. The top of the chamber was sealed with a coverslip and imaged at 63x magnification with a water-immersion objective using a Zeiss LSM 800 microscope. The images were cropped and processed in FIJI to remove background fluorescence (rolling ball radius of 50 pixels), and increase brightness, sharpness, and contrast.

### Synthetic Mucus (SM) Hydrogel Preparation

A previously established protocol was adapted to prepare synthetic mucus hydrogels with and without sNETs and its individual components.^32^ Briefly, porcine gastric mucins (PGM; Sigma Aldrich; mucin from porcine stomach, type III, bound sialic acid 0.5-1.5%, partially purified powder) were stirred for 2 hours at a concentration of 40 mg/ml in the same physiological buffer used to make the sNETs. Solutions of 3 mg/ml DNA, histones, sNETs, or PGM were prepared separately using the same buffer type. 4 arm-PEG-thiol (PEG-4SH; Laysan Bio) was used as a crosslinker for the mucins. The PEG-4SH was added directly into each of the 3 mg/ml DNA, histones, sNETs, or additional PGM solutions at a concentration of 20 mg/ml. The 40 mg/ml PGM solution and 20 mg/ml PEG-4SH crosslinking solution containing 3 mg/ml DNA, histones, sNETs, or additional mucins were combined in equal volumes and mixed using a pipette.

### Macrorheology

SM hydrogels with and without sNETs (n = 3 per condition) were cast in molds with a 20 mm diameter at a depth of ∼2 mm and gelled at room temperature for 48 hours. The viscoelastic properties of SM hydrogels were measured using an ARES G2 rheometer (TA instruments) with a 20 mm parallel plate geometry. A strain sweep was performed between 0.1 and 10% strain to determine the linear viscoelastic region at an angular frequency (ω) of 1 rad/s. Subsequently, a frequency sweep between 0.1 and 100 rad/s was conducted within the linear viscoelastic region of the gel at 1% strain to determine the elastic (G’(ω)) and viscous moduli (G’’(ω)).

### Preparation of Nanoparticles for Microrheology

A surface coating of polyethylene glycol (PEG) was added to 100 nm carboxylate modified red fluorescent polystyrene (PS) nanoparticles (Life Technologies). 5 kDa methoxy PEG-amine (Creative PEGworks) was attached to the surface of the PS nanoparticles by a carboxyl-amine linkage, as previously described.^32,35^ In order to confirm attachment of the PEG to the surface, the zeta potential of the particles was measured with a Nanobrook Omni Particle Analyzer (Brookhaven Instruments).

### Multiple Particle Tracking Microrheology

For microrheology experiments, 25 μl of hydrogel was added to a microscopy chamber created by sticking a vacuum-grease covered O-ring to a glass microscope slide. 1 μl of PEG-coated PS nanoparticles diluted to 0.0025% w/v in ultrapure water was added to the hydrogel solution, then the microscopy chamber was sealed using a cover slip. The hydrogels (n = 3 per condition) gelled for 24 hours at room temperature in the sealed microscopy chamber. For experiments where DNase was added to the hydrogels, 24 μl of hydrogel solution was added to the microscopy chamber. 1 μl of the PS-PEG nanoparticles diluted to 0.0025% w/v in ultrapure water was added, then the hydrogels (n = 3 per condition) were gelled for 22 hours at room temperature with no coverslip covering the microscopy chamber well. Instead, the microscope slides were sealed inside humidified containers using parafilm. DNase was dissolved in the physiological buffer following 22 hours of gelation. The solution of DNase was added to the surface of the hydrogels at a final concentration of 7 μg/ml and coverslips were added to seal the microscopy chamber. All hydrogels were incubated at 37°C for 1 hour, then imaged using fluorescence video microscopy. Nanoparticle diffusion was imaged using 63x magnification with a water-immersion objective on a Zeiss LSM 800 microscope. 10 second videos of the fluorescent nanoparticles were captured within each hydrogel at a frame rate of 33 Hz in 5 different regions of the sample. Using the videos of the 100 nm fluorescent PS-PEG nanoparticles, the MSD as a function of the lag time (τ) was quantified for each particle in MATLAB using the equation ⟨Δ*r*^2^(τ)⟩ = ⟨(*x*^2^ + *y*^2^)⟩. Once the MSD values were obtained, the generalized Stokes Einstein equation was used to calculate the viscoelastic spectrum 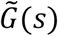 with the Laplace transform of ⟨Δ*r*^2^(τ)⟩, which is ⟨Δ*r*^2^(*s*)⟩. The viscoelastic spectrum was calculated with the equation 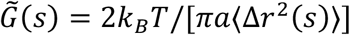 where *k*_b_*T* is thermal energy, *a* is particle radius, and *s* is complex Laplace frequency. Then, the complex modulus of the gel can be computed using the equation *G*^∗^(*ω*) = *G′(ω)* + *G″* (*iω*). The variable *s* is replaced by *iω*, where *i* is a complex number and *ω* is the frequency. The pore size of the hydrogel (ξ) can then be calculated using *G*’ with the equation 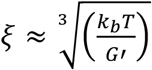.^28^

### HAE Culture

The immortalized airway basal cell line BCI-NS1.1 (basal cells from non-smoker) was grown in an incubator set to 37°C and 5% CO_2_. The BCI-NS1.1 cells were initially grown to ∼70-80% confluence in a tissue culture flask using Pneumacult Ex Plus Media (Stem Cell Technologies). Cells were then seeded on Millicell cell culture inserts (Millipore Sigma; 0.4 μm pore PET membrane) in a 24 well plate at a density of ∼10,000 cells/cm^2^. The BCI-NS1.1 cells on the culture inserts were initially submerged in Pneumacult Ex Plus media, which was added to both the apical and basolateral compartments until cells were a confluent monolayer. Once confluent, the media in the apical compartment was removed and the cells were grown at the air-liquid interface (ALI). While growing at ALI, the media in the basolateral compartment was changed to Pneumacult ALI media (Stem Cell Technologies). Cells were grown at ALI for at least 28 days to differentiate into the various cell types found in the airway. Media in the basolateral chamber was replaced every other day. After 21 days at ALI, mucus was washed from the surface of the cultures 2 times per week. Mucus was washed from the surface of each culture by applying sterile PBS to the apical surface. After 30 minutes of incubation (37°C and 5% CO_2_), the PBS was removed and the washings could be collected to isolate the mucus. Mucus was isolated from collected PBS washings using 100 kDa Amicon Ultra centrifuge filters (Millipore Sigma). The washings were centrifuged for 20 minutes at 14,000 g to remove the PBS and isolate mucus secreted by the cells.

### TEER

The TEER of the cultures was measured using a Millicell Electrical Resistance System ERS-2 Volt-Ohm Meter (Millipore). Fully differentiated BCI-NS1.1 cultures (n = 3) were washed immediately prior to the first TEER reading, and the collected mucus was pooled and aliquoted. The probe was sterilized in 70% ethanol for 15 minutes, then washed using PBS (no Ca^2+^ or Mg). The media was removed from the basolateral compartment of the cells at ALI, and PBS was added to both the apical and basolateral compartments and cells were left to come to room temperature. TEER measurements were conducted using the probe, including a blank cell culture insert. The final TEER measurement was determined by subtracting the TEER value of the blank insert from the TEER value of each culture insert with cells growing, then multiplying by the area of the insert (0.3 cm^2^). Once the first measurement had been completed, PBS was removed and media was added back to the basolateral compartment. Then, sNETs were combined in an equal volume ratio with the freshly collected mucus aliquots and 6 μl was transplanted back onto the apical surface of the cultures. Cells with the transplanted mucus were placed back into the incubator for 24 hours, then TEER was measured again. The transplanted mucus was not removed from the surface of the cultures to read the TEER after 24 hours.

### Mucociliary Transport and Cilia Beat Frequency

Mucus was washed from the surface of each HAE culture prior to the experiment using PBS and isolated using 100kDa centrifugal filters. All mucus was pooled together, then two different aliquots were made. An equal volume of sNETs suspended in the physiological buffer was added to the HAE mucus aliquot and mixed thoroughly (sNET+ HAE mucus). The sNETs were prepared at 3 mg/ml, and diluted to 1.5 mg/ml final concentration when mixed with the HAE mucus. As a vehicle control, an equal volume of the physiological buffer alone (the same buffer that was used to make the sNET suspension) was mixed into the second remaining HAE mucus aliquot. A volume of 6 μl of mucus containing either the buffer alone (vehicle) or 3 mg/ml sNETs was transplanted onto the surface of each culture (n = 3 per condition), making sure that the mucus covered the entire apical area of the culture. Then, 2 μm fluorescent polystyrene microspheres (Life Technologies) were sonicated for 7 minutes and diluted 1:3000 in sterile PBS. A volume of 5 μl of the diluted microspheres was pipetted onto the apical surface of each culture with the transplanted mucus already applied. Cultures were then incubated for 30 minutes at 37°C before imaging. Cultures were imaged using a Zeiss 800 LSM microscope at 10x magnification. Videos of the beads moving on the apical surface of the mucus layer were taken at a frame rate of 0.5 Hz for 10 seconds in 3 different regions for each culture. Tracking of the microspheres was performed using a custom MATLAB algorithm to analyze each frame of the video and quantify the trajectories of the microspheres in the *xy* plane. The rate of mucociliary transport was calculated by dividing the displacement of each microsphere by the total time. In addition, 10 second videos of the culture surface taken using 10x magnification at a frame rate of 20 Hz in bright field were recorded in 3 different regions of each culture. The ciliary beat frequency was quantified using a custom MATLAB algorithm where the number of local pixel intensity maxima is counted to indicate the beating of cilia. The ciliary beat frequency was then determined by quantifying the number of cilia beats over the total time.

For the DNase experiment, cultures (n = 3 per condition) were first imaged pre-treatment with only the transplanted mucus and beads applied to the surface. Immediately afterwards, DNase was applied to the apical surface at a final concentration of 7 μg/ml, and cultures were incubated for another 30 minutes. Cultures were then imaged again to track the movement of the beads after the application of DNase. Videos of the movement of the beads before and after DNase application were taken using the 10x objective of the Zeiss 800 LSM microscope at a frame rate of 0.5 Hz for 20 seconds in 3 different regions for each culture. The analysis process was the same as described above to determine the mucociliary transport rate. Cilia beat frequency was determined using the same aforementioned imaging and analysis methods.

### Bacterial Culture

Planktonic green fluorescent protein (GFP) expressing *Pseudomonas aeruginosa* (PAO1) bacteria (ATCC 10145GFP) were grown in Lennox Broth (LB) media using a shaking incubator at 37°C. The optical density at 600 nm (OD_600_) of the culture was measured immediately before use with a BioTek plate reader, and diluted to the desired OD_600_ using the physiological buffer.

### Fluorescence Microscopy of Bacteria in HAE Mucus

Mucus was washed from the surface of each HAE culture prior to the experiment using PBS and isolated using 100kDa centrifugal filters. All mucus was pooled together, then two different aliquots were made. An equal volume of the sNET suspension was added to the mucus aliquot and mixed thoroughly. As a vehicle control, an equal volume of physiological buffer was mixed into the second remaining mucus aliquot. 25 μl of each sample of mucus was added to a microscopy chamber created using an O-ring adhered to a glass slide with vacuum grease. 1 μl of PAO1 diluted to an OD_600_ of 0.01 in the physiological buffer was added to the mucus samples in the microscopy chamber. The top of the chamber was then sealed with a cover slip and the slide was placed in an incubator at 37°C. After 24 hours, the slides were imaged using 10x magnification with a Zeiss LSM 800 microscope. Images were processed in FIJI, where the background fluorescence was subtracted (rolling ball radius of 50 pixels) and brightness and contrast were increased.

### Fluorescence Readings of Bacteria in HAE Mucus

HAE mucus was collected and isolated as described above from fully differentiated BCI-NS1.1 cultures. Immediately following isolation, the mucus was frozen at -80°C for long-term storage. Mucus was thawed overnight at 4°C, then pooled together. 50 μl of mucus was added to the wells (n = 3 per condition) of a half area black flat bottom plate (Corning). PAO1 cultures grown in LB media were diluted to an OD_600_ of 0.1 in buffer, then 1 μl (10^4^ bacterial cells) was added to each well containing the mucus. The plate was covered with a breathe-easy sealing membrane (Diversified Biotech) and incubated overnight at 37°C. A plate reader (BioTek) was used to read the GFP fluorescence of the wells after 24 hours to obtain a baseline fluorescence level. After the initial fluorescence reading at 0 hours, 50 μl of either the sNET suspension or buffer alone was incorporated into the mucus by mixing up and down with a high-viscosity pipette. The sNETs were prepared at 3 mg/ml, and diluted to 1.5 mg/ml final concentration when mixed with the mucus. The plates were covered with a breathe-easy sealing membrane and incubated at 37°C. Fluorescence readings of GFP were taken at 3, 6, and 24 hours after incorporation of sNETs or buffer to indicate bacterial viability. Blank wells with no bacteria added for each condition were included and their fluorescence values were subtracted after each reading to account for background fluorescence.

## Results and Discussion

### Characterization of sNETs

To formulate sNETs, DNA and histones were dissolved and combined in equal ratios to cause complexation via electrostatic interactions, with DNA being negatively charged and histones being positively charged. Previous work found that using equal ratios of 50 wt% DNA and 50 wt% histones to synthesize sNETs produced the most structural similarity to endogenous NETs.^26,30^ To create a suspension of sNETs, the large DNA-histone complexes were broken up using ultrasonication (**Figure 1A**).^26,30,31^ Fluorescence microscopy of the sNETs revealed successful recreation of a porous DNA scaffold structure, similar in appearance to endogenous NETs (**Figure 1B**).^8^ We found the DNA network of the sNET can trap and sterically confine the diffusion of 100 nm nanoparticles with a dense surface coating of PEG, indicating similar structure and functionality to NETs in trapping microbes in the web-like matrix (**Video S1**). To estimate the conversion of free DNA and histones into sNETs quantitatively, a suspension of sNETs made at a concentration of 3 mg/ml was centrifuged to pellet the sNETs, then the remaining free DNA concentration in the supernatant was measured to be 0.15 mg/ml (**Table 1**). We estimated ∼95% conversion of DNA and histones into sNETs using this procedure (**Table 1**). The A260/280 ratio of the DNA found in the supernatant of the sNET suspension was 1.83, suggesting that the remaining DNA in the supernatant was purely free DNA and not in complexation with histones.^36^ When compared to the same assay done on a solution of dissolved DNA at 3 mg/ml, the free DNA content in the supernatant was 2.73 mg/ml with an A260/280 ratio of 1.84. Finally, the zeta potential of the sNETs in suspension was measured as -4.75 ± 1.67 mV, near neutral in charge due to the electrostatic interactions of the DNA and histones (**Table 1**).

**Table 1.** Characterization of sNETs. Efficiency of DNA-histone complexation to form sNETs is estimated based on quantification of free DNA and measurement of zeta potential of sNETs.

| Initial Concentration of sNETs (mg/ml) | Concentration of DNA in Supernatant (mg/ml) | Approximate % Conversion to sNETs | Zeta Potential (mV) |
| --- | --- | --- | --- |
| 3 | 0.15 | 95% | $-4.75 \pm 1.67$ |

**Figure 1.**
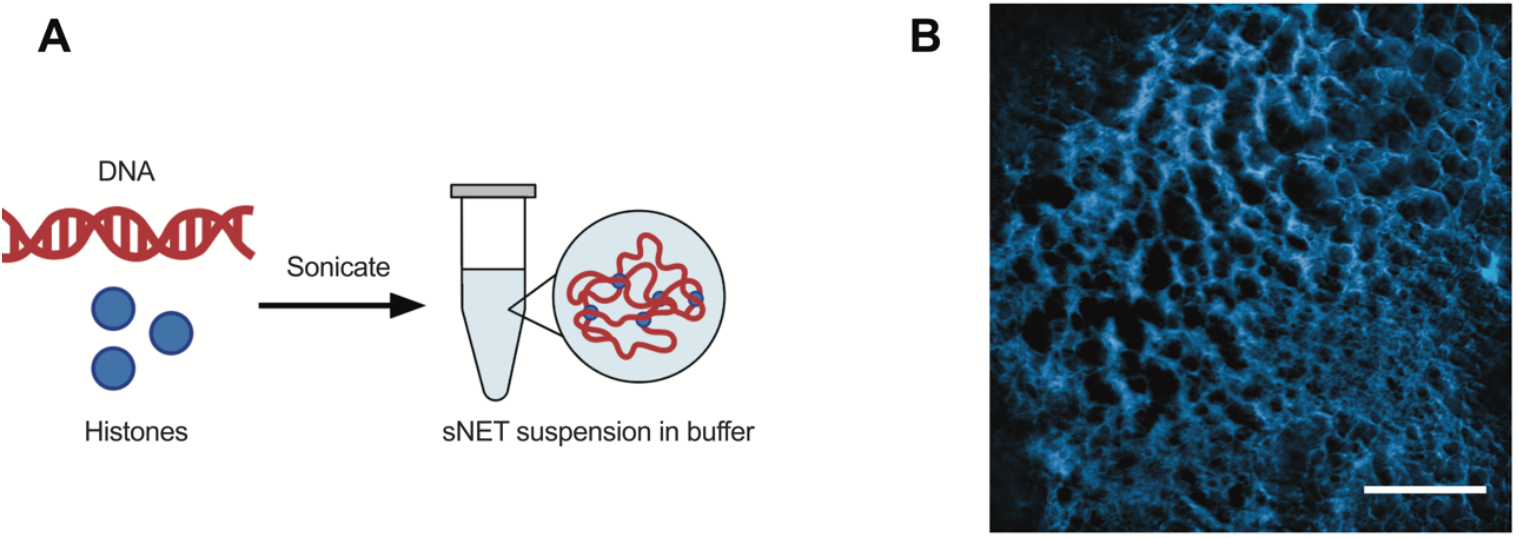
Synthetic NET (sNET) formulation. (A) Schematic of sNET formation by combining 50 wt% DNA and 50 wt% histones dissolved separately in buffer. The DNA and histone will electrostatically interact to form large complexes that can be sonicated to create a suspension of sNETs. (B) Fluorescence microscopy image of the DNA structure of an sNET, stained with DAPI. Scale bar, 25 μm.

### sNETs Alter Synthetic Mucus Microstructure and Viscoelasticity

The primary solid component of mucus are mucin proteins that crosslink via disulfide bonds to form a viscoelastic gel. To determine the effect of NETs on the viscoelastic properties and microstructure of mucus, sNETs were incorporated into a synthetic mucus (SM) hydrogel. Use of the SM model enabled us to precisely control the composition and total percentage solids concentration within sNET–SM hydrogel. The SM hydrogel model was developed in our previous work using PEG-4SH to form disulfide bonds between the mucins.^32–34^ SM was prepared by combining 4% PGM solution with a solution containing 2% PEG-4SH and either 0.3% sNET, free DNA, free histones, or additional PGM (**Figure 2A**). The resulting normal (PGM only) or CF-like (sNET containing) SM hydrogels had the same total percent solids concentrations. The final concentration of sNET was based upon reported concentrations of extracellular DNA in CF sputum samples, which span a wide range of up to ∼10 mg/ml but average around 3 mg/ml.^27,28,37–39^ In contrast, one study reported healthy human mucus contains an average extracellular DNA concentration of 0.006 ± 0.003 mg/ml.^37^ Previous studies have found that approximately 50% of the measured extracellular DNA in CF sputum is in complexation with proteins indicating that it is a NET, therefore the final concentration of sNET used in this study was 1.5 mg/ml (0.15% w/v).^41,42^ Other extracellular DNA sources may include that from necrotic host cells, especially neutrophils, not undergoing NETosis.^27^

**Figure 2.**
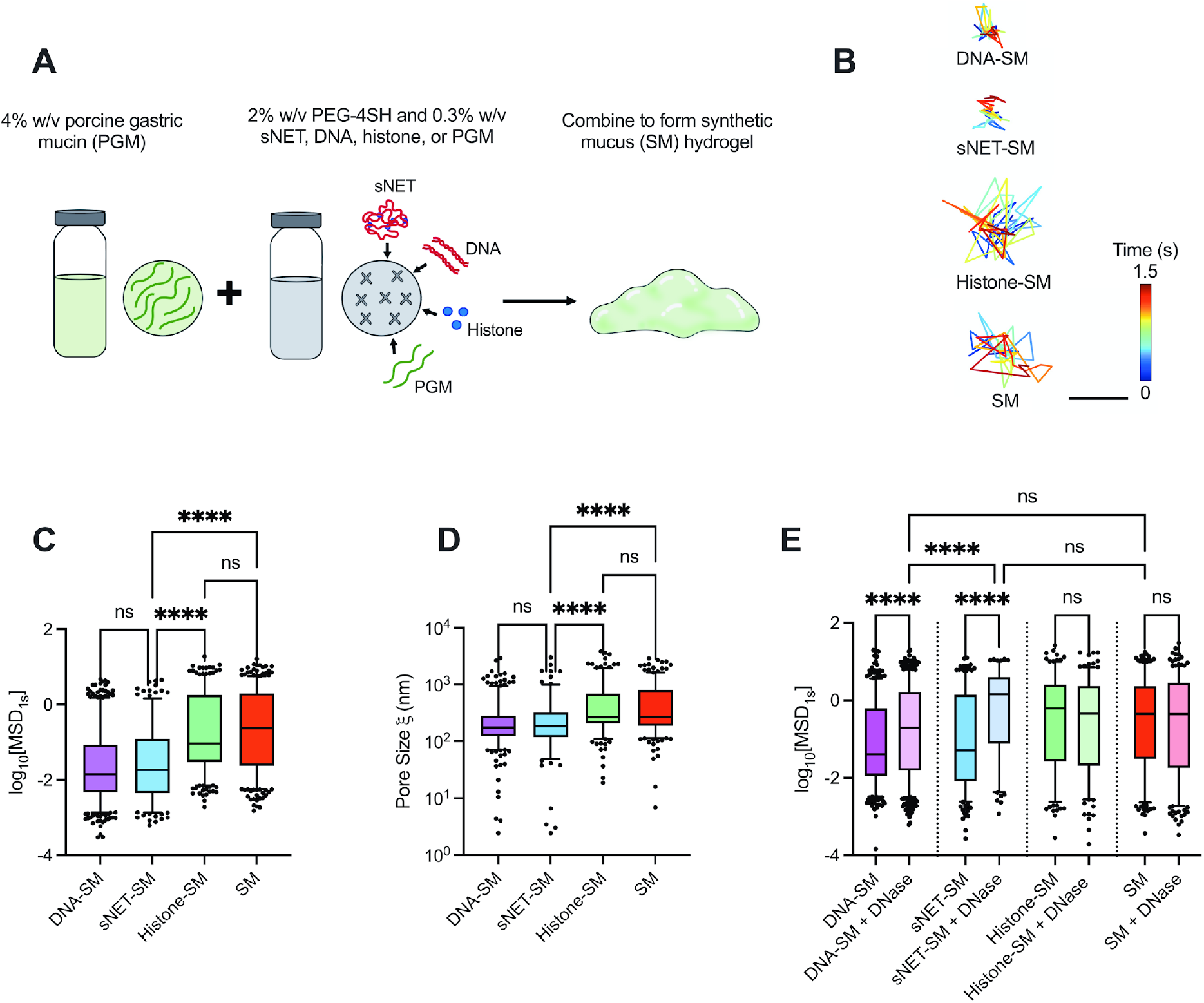
sNETs Alter Synthetic Mucus Microstructure. (A) Schematic of synthetic mucus hydrogel preparation with various additives (sNETs, DNA, Histones, or additional PGM). (B) Representative trajectories of PEGylated 100 nm nanoparticle diffusing within each hydrogel type for 1.5 seconds. Scale bar, 200 nm. Box and whisker plots of (C) the logarithm based 10 of the measured MSD at 1 second (log_10_[MSD_1s_]) of PEGylated 100 nm nanoparticles in each hydrogel type (n = 3 replicates per condition, with 5 different randomly chosen regions of each individual gel imaged), and (D) the estimated pore size of each hydrogel type calculated based on measured MSD. (E) Box and whisker plot of the log_10_[MSD_1s_] of 100 nm PEGylated nanoparticles measured for each hydrogel type with and without 7 μg/ml DNase I treatment for 1 hour at 37°C (n = 3 replicates per condition, with 5 different randomly chosen regions of each individual gel imaged). For (C–E), statistical significance determined by Kruskall Wallis test with Dunn’s multiple comparisons test (**** = p < 0.0001).

We performed multiple particle tracking analysis on each of the SM types using 100 nm muco-inert nanoparticles to evaluate the microrheological and microstructural properties of SM containing sNETs and sNET components. **Figure 2B** shows representative trajectories of a single nanoparticle diffusing in each hydrogel type. The nanoparticle diffusion rate was assessed quantitatively by determining the mean squared displacement (MSD). The hydrogels containing free DNA (DNA-SM) and sNETs (sNET-SM) showed a significant reduction in measured MSD in comparison to the hydrogels containing histones (Histone-SM) and PGM only (SM) (**Figure 2C**). The measured MSD of each nanoparticle was also used to calculate the approximate pore sizes (*ξ*) of each hydrogel type. We found smaller pore sizes in the hydrogels containing sNETs or DNA compared to those containing histones or PGM only (**Figure 2D**). These data are consistent with a previous study finding similar results with endogenous NETs reducing mucus network pore size.^25^ Our data indicate that the DNA component of the sNETs is likely responsible for the changes to SM microstructure. Extracellular DNA, including that which forms NETs, has been previously described as an important macromolecule contributing to enhanced CF mucus viscoelasticity.^6,25,28,43^

DNase is a clinically used inhaled mucolytic agent used by a large proportion of CF patients to degrade extracellular DNA, including NETs, present in airway mucus.^38^ Therefore, it was important to determine if our model containing sNETs were responsive to DNase. In order to determine how DNase affected the microstructure, we treated each hydrogel type with DNase for 1 hour at a final concentration of 7 μg/ml, which is the reported concentration found in the sputum of CF patients after nebulization.^28,44^ The MSDs of the nanoparticles increased significantly in the hydrogels containing DNA and sNETs when treated with DNase, and there was no significant change in the MSDs of nanoparticles in the hydrogels containing histones or PGM only with DNase treatment (**Figure 2E**). The MSDs of the nanoparticles in the hydrogels with DNA and sNETs treated with DNase were increased to be very similar to the PGM only hydrogel.

In addition to assessing microstructural changes, we evaluated the viscoelastic properties of the PGM only and sNET containing hydrogels using macrorheology. The formulation of both hydrogels remained the same as the microrheology, and were therefore the same total percentage of solids. We found that the elastic (G’) and viscous (G’’) moduli and the complex viscosity (*η*^∗^) were increased in the hydrogel containing sNETs (**Figure 3A-C**). The rheological properties for SM and sNET-SM are similar in magnitude to human mucus from healthy individuals and individuals with CF, respectively.^45,46^ Our microrheology results indicated that incorporating sNETs into our SM hydrogel model caused changes to the microstructure, namely reduction in pore size, that could be reversed with DNase application. Though the pore sizes were estimated based on the MSD, we can still observe a trend in a tightened network and smaller pore size with the addition of sNETs to the SM hydrogel. Macrorheology revealed that sNETs increase viscoelastic moduli and complex viscosity of the SM hydrogel. This supports that the addition of sNETs to our mucin hydrogel is able to recapitulate the effects of NETs in CF mucus in tightening the microstructural network formed by mucins and subsequently increasing mucus viscoelasticity.

**Figure 3.**
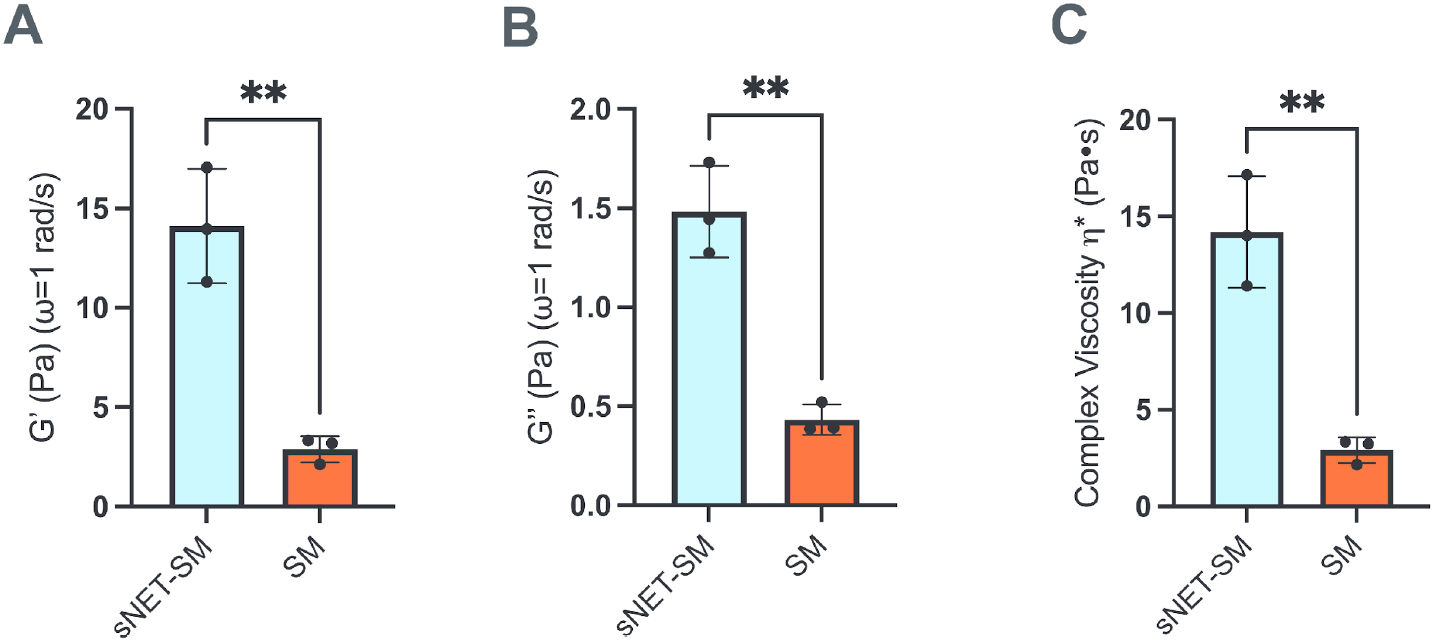
sNETs Increase Synthetic Mucus Viscoelasticity. (**A**) Mean elastic, (**B**) viscous moduli (*G*′, *G*″) and (**C**) complex viscosity (η*) at ω = 1 rad/s for SM and sNET-SM hydrogels (n = 3 replicates per condition). Statistical significance determined by unpaired two tailed t-test (** = p < 0.01).

### sNETs Decrease Mucociliary Transport *In Vitro*

To examine the impact of sNET on mucociliary clearance, we next constructed an *in vitro* model using the normal human airway epithelial (HAE) immortalized cell line BCI-NS1.1, capable of producing mucus and performing MCT. We washed the secreted HAE mucus off the apical surface of fully differentiated cultures grown at air-liquid interface (ALI) and pooled the harvested HAE mucus. A suspension of sNETs prepared at 3 mg/ml was added to the HAE mucus and thoroughly incorporated (sNET+) for a final concentration of 1.5 mg/ml sNETs. In addition, HAE mucus containing the same buffer that the sNETs were suspended in was used as a vehicle control (control). We overlaid the sNET+ or control HAE mucus onto the apical surface of the freshly washed cultures, then measured the rate of MCT. Fluorescent 2 μm microspheres were added to the apical surface atop the newly overlaid sNET+ or control HAE mucus. We then tracked the movement of these microspheres across the mucosal surface of the cultures to measure MCT rate (**Figure 4A**). We found MCT rate was significantly reduced in the sNET+ HAE mucus compared to the control HAE mucus, with no changes in cilia beat frequency (CBF) (**Figure 4B-C**). Cultures with sNET+ HAE mucus overlaid display very little transport of the microspheres in comparison with cultures with the control HAE mucus overlaid (**Figure 4D**). We also confirmed the addition of the sNET+ HAE mucus to the apical surface of the cultures did not affect the transepithelial electrical resistance (TEER), indicating that the transfer had little impact on the cell viability or monolayer integrity (**Figure S1**).

**Figure 4.**
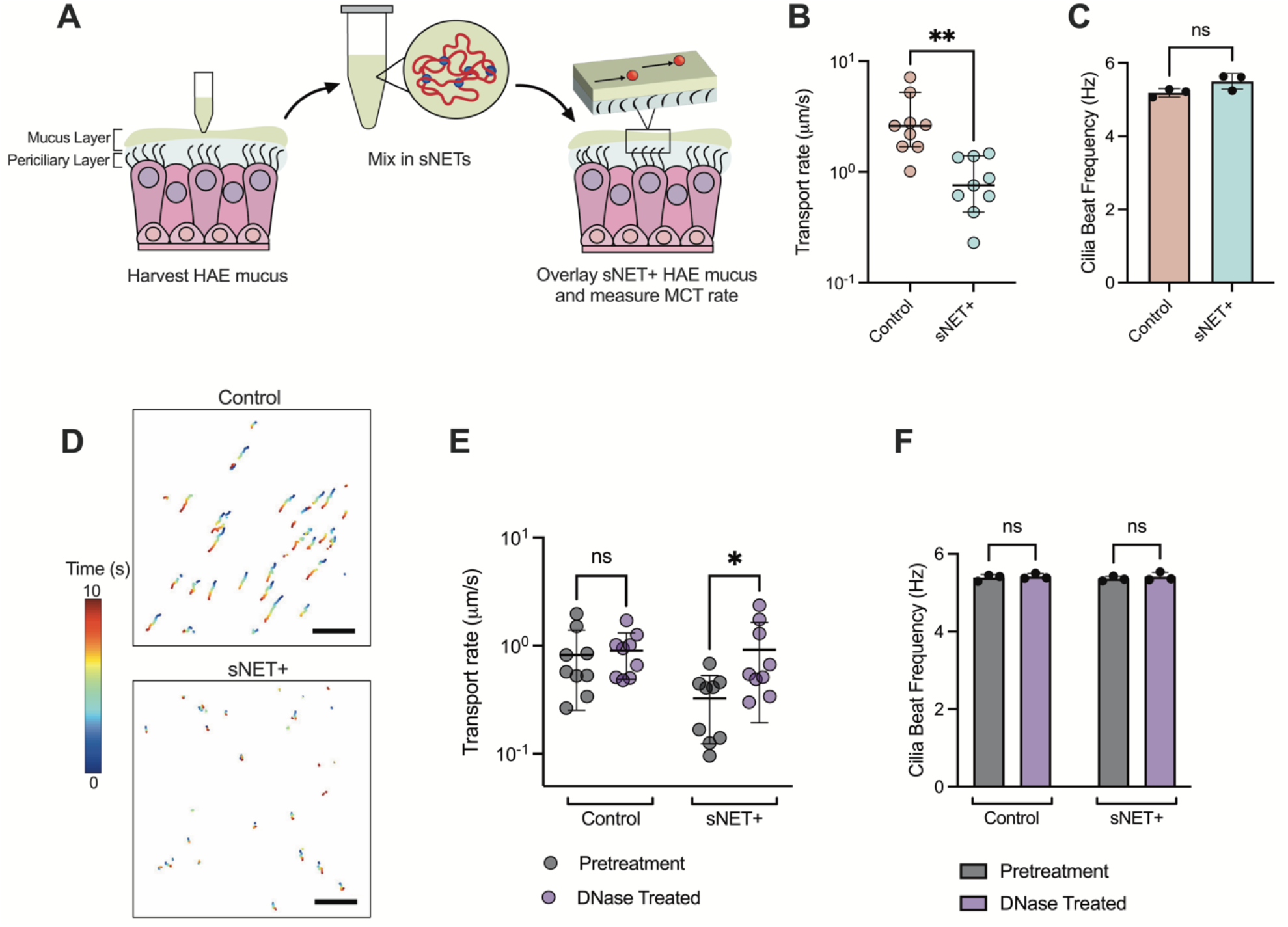
sNETs Decrease Mucociliary Transport *In Vitro*. (A) Schematic of the process of washing cultures to harvest HAE mucus, then incorporating sNETs into the fresh HAE mucus (sNET+ HAE mucus), and overlaying the sNET+ HAE mucus back onto the surface of the cultures to measure MCT using fluorescent microspheres. (B) Median transport rates of the 2 μm microspheres and (C) the mean cilia beat frequency of cultures overlaid with control and sNET+ HAE mucus (n = 3 replicates per condition, with 3 randomly chosen regions of each individual culture imaged). (D) Representative trajectories of the microspheres being transported across the mucosal surface of cultures over the course of 10 seconds with control or sNET+ HAE mucus applied. Scale bar, 100 μm. (E) Median transport rates and (F) mean cilia beat frequency measured from the same set of cultures with either control or sNET+ HAE mucus overlaid before (pretreatment) and after (DNase treated) application of DNase I at a concentration of 7 μg/ml for 30 minutes at 37°C (n = 3 replicates per condition, with 3 randomly chosen regions of each individual culture imaged). Statistical significance determined by unpaired two tailed t-test for (B, C), a two-way repeated measures ANOVA test for (E, F) (Ns = p > 0.05, * = p < 0.05).

In addition, we measured the MCT rate and CBF of cultures with either sNET+ or control HAE mucus overlaid both before and after DNase treatment. After an initial pretreatment measurement of the transport rate and CBF, DNase was applied at a final concentration of 7 μg/ml for 30 minutes and both the transport rate and CBF were measured again. Our results show that there was no change in the MCT rate of the control HAE mucus after DNase treatment, but that the transport rate became faster after the application of DNase to the cultures with sNET+ HAE mucus, with no changes in CBF before or after treatment (**Figure 4E-F**). Our *in vitro* model shows the ability of sNETs to mimic the decrease in MCT seen in CF as a result of excess NETs that enhances the viscosity of the mucus layer (**Figure 3**). We note our studies here were performed with an immortalized non-CF HAE cell line. However, the methods employed here could be easily adopted using primary CF HAE cells in future work.

### sNETs Alter Bacterial Growth Patterns and Sustain Survival in Mucus

As previously discussed, many CF patients experience chronic infection of the airways with *P. aeruginosa*. Therefore, we characterized the interactions of the planktonic green-fluorescent protein (GFP) expressing strain of *P. aeruginosa* bacteria, PAO1, with sNET+ HAE mucus in comparison to control HAE mucus. After growing for 24 hours in control HAE mucus, the bacteria formed very small aggregate colonies (**Figure 5A**). However, 24 hours after inoculation in sNET+ HAE mucus, the bacteria were localized to much larger areas, presumably due to their capture by the sNETs (**Figure 5B**). A similar phenomenon of bacterial localization to the sNETs was observed when bacteria were added to the suspension of 1.5 mg/ml sNETs alone in the absence of mucus and incubated for an hour before imaging (**Figure S2**). Next, we grew PAO1 in HAE mucus, then subsequently added sNETs into the HAE mucus to simulate the innate immune response of neutrophils migrating to the site of infection and releasing NETs. We measured the GFP fluorescence of the bacteria as an indicator of viability at 0 h immediately before adding either a suspension of sNETs at a final concentration of 1.5 mg/ml (sNET+ HAE mucus) or an equal volume of the buffer that the sNETs were prepared in as a vehicle control to the HAE mucus. Fluorescence readings were repeated at 3, 6, and 24 h after introducing sNETs or the buffer alone. We found that the GFP fluorescence steadily declined over time in the control HAE mucus (**Figure 5C**). However, the addition of the sNETs to the HAE mucus sustained the survival of the bacteria over time as indicated by the fluorescence readings (**Figure 5D**).

**Figure 5.**
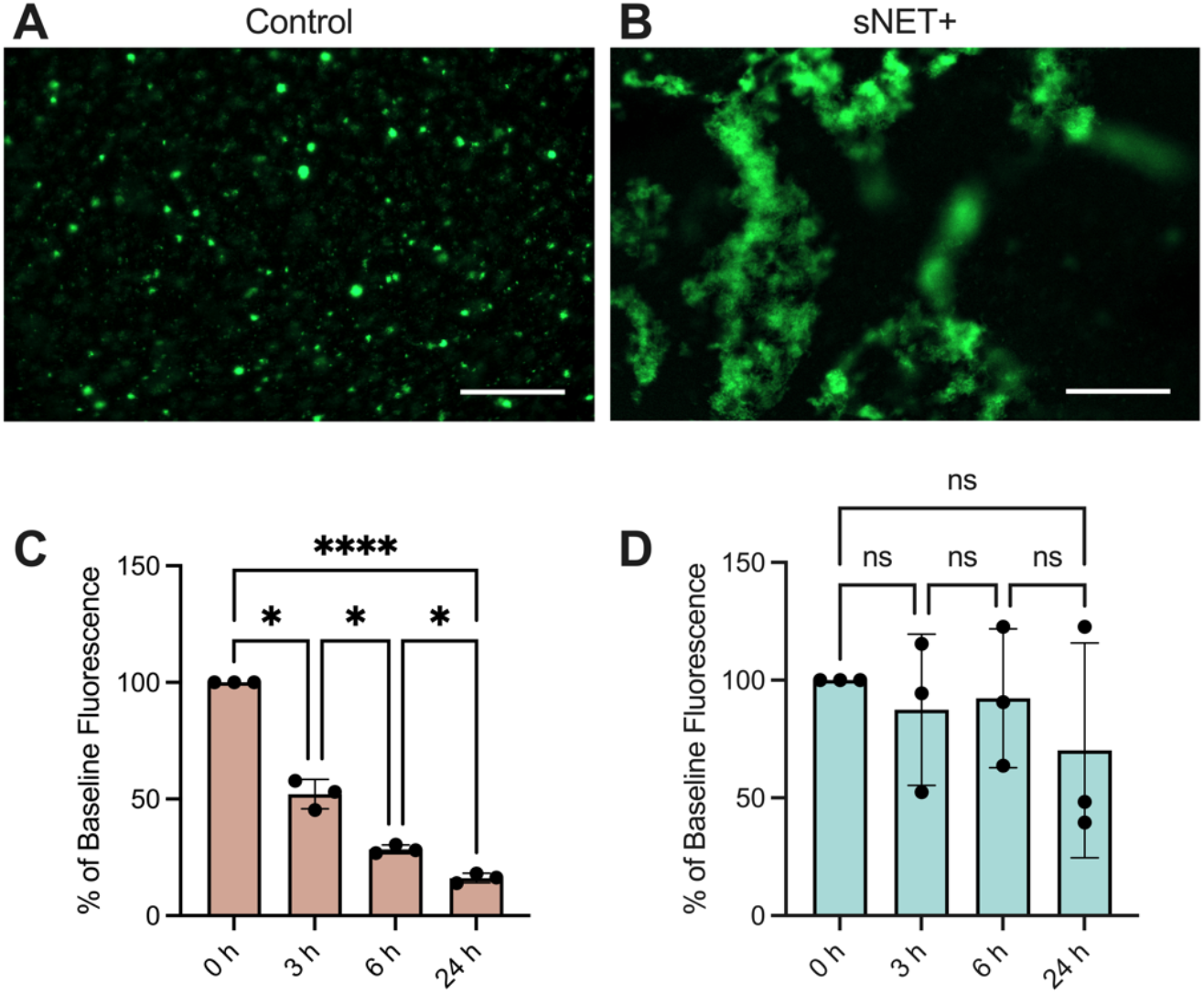
sNETs Alter Bacterial Growth Patterns and Sustain Survival in Mucus. Fluorescence microscopy images of GFP-expressing PAO1 (GFP PAO1) bacteria grown for 24 hours at 37°C in (A) control or (B) sNET+ HAE mucus. Scale bar, 100 μm. GFP PAO1 were grown in (C) control or (D) sNET+ HAE mucus for 24 hours and the GFP fluorescence of the bacteria was measured at 0, 3, 6, and 24 hours (n = 3 replicates per condition). Values are plotted as percentage of the baseline fluorescence at 0 h with background fluorescence subtracted from wells containing either control or sNET+ HAE mucus with no bacteria. Statistical significance determined by one-way repeated measures ANOVA tests with Tukey’s multiple comparisons tests (Ns = p > 0.05, * = p < 0.05, **** = p < 0.0001).

Our results align with previous work by Song et al that found that formulations of the sNET material alone containing an equal ratio of DNA and histones at concentrations exceeding 0.5 mg/ml promoted bacterial survival and biofilm formation by *Staphylococcus aureus*, another bacterial species commonly found in CF airways.^30^ The proposed explanation was that the electrostatic interactions between the sNET material and bacterial membranes encouraged bacterial adhesion and aggregation.^30^ We similarly observed *P. aeruginosa* forming large aggregates presumably through their interactions with sNETs within mucus, corroborating previous reports that NET-mediated killing of the bacteria is impaired and evidence that NETs are likely beneficial in establishing biofilms in the CF mucosal environment.^12,15^ Taken together, our results demonstrate that the addition of sNETs in mucus alters bacterial growth patterns, enhances survival, and ultimately better simulates the CF airway where bacteria can flourish and cause chronic infection. Additional work examining the long-term impacts (i.e. >24 hours) of sNETs on bacterial survival, response to antibiotics, and other patho-adaptations of *P. aeruginosa* observed in CF^47,48^ are needed to further confirm the relevance of these models to chronic lung infections. We are also aware endogenous NETs contain other species (e.g. antimicrobial peptides) which could influence growth behavior of *P. aeruginosa*. It should be noted sNETs prepared with other NET-associated proteins has been recently reported^31^ where we plan to adapt these procedures for future studies.

## Conclusions

In this work, we established new biomaterial-based *in vitro* models capable of recapitulating the impacts of NETs on critical aspects of CF lung disease including mucus hyperviscosity, impaired mucociliary transport, and bacterial colonization. Given the well-established role of innate immune dysfunction and NETosis in CF, it is important to consider the impact of NETs in preclinical CF models used in development of new therapies and studies on host-pathogen interactions. Our results establish a highly valuable means to incorporate sNET biomaterials into *in vitro* lung airway models to better reflect the CF lung microenvironment. Inhaled DNase therapy is frequently prescribed to CF patients^5^ and it should be noted that traditional *in vitro* models are unresponsive to this treatment. Importantly, we found our *in vitro* model using sNET biomaterials responds to DNase treatment with improvements in MCT at a physiologically relevant dose. As noted, NETs are comprised of additional components such as neutrophil elastase, alpha-1 antitrypsin, and antimicrobial peptides.^12^ Future work incorporating these components into sNETs for use in this *in vitro* CF model will help to further elucidate their individual and collective contributions to CF lung disease progression.

## Author Contributions

A.B. conceived, designed, and performed experiments. S.Y. designed experiments and assisted with bacterial culture. G.A.D. conceived and designed experiments. A.B. and G.A.D. wrote the article. All authors reviewed and edited the article.

## Acknowledgements

This work was supported by the Burroughs Wellcome Fund Career Award at the Scientific Interface (to G.A.D.), Cystic Fibrosis Foundation (BOBOLT23H0 to A.B.), the National Science Foundation (DGE1840340 to S.Y., 2129624 to G.A.D.) and the National Institutes of Health (R21 EB030834, R01 HL160540 to G.A.D.).

