## Supplemental Information for "Engineering *in vitro* models of cystic fibrosis lung disease using neutrophil extracellular trap inspired biomaterials"

**Description for Video S1.** 100 nm PEG-coated nanoparticles diffusing within a DAPI-stained DNA scaffold of sNETs in suspension. Video captured at 63x magnification with a Zeiss 800 LSM microscope, taken at a frame rate of 0.65 Hz. The original video (in color) is shown on top, then the nanoparticle trajectories were tracked using the FIJI, shown on the bottom. The video was converted to a stack of 8-bit images to be compatible for use with the FIJI trackmate plugin. Nanoparticles (circled in red) were tracked, with their trajectory paths shown in white. Nanoparticles with no white trajectory paths shown indicate that the particle was immobilized. Scale bar, 25 µm. The video is looped to play 5 times through.

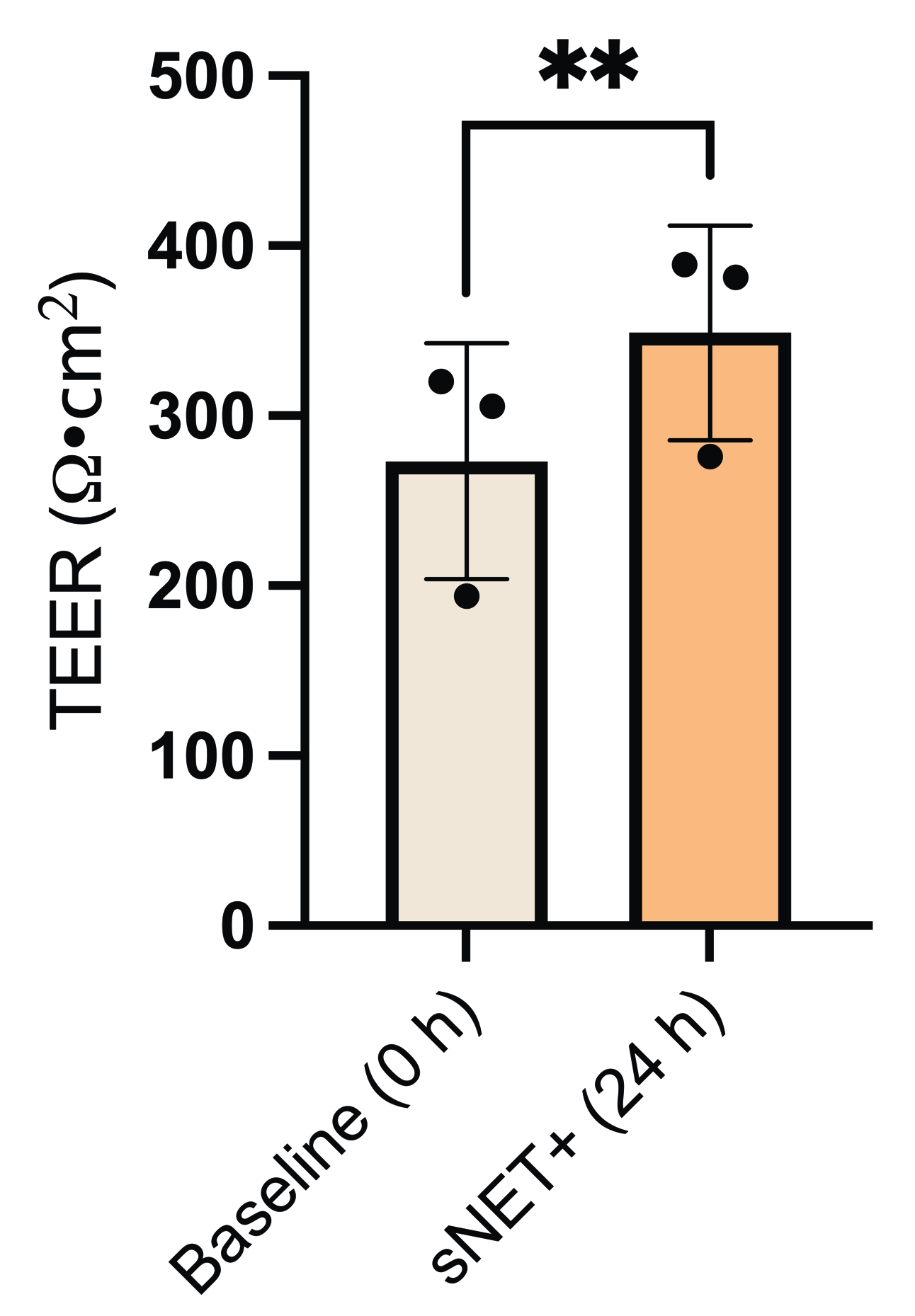

**Figure S1.**  Mean TEER of BCI-NS1.1 cultures (n = 3) at air-liquid interface measured immediately before (baseline, 0 h) and 24 hours after overlaying sNET+ HAE mucus onto the apical surface. Statistical significance was determined using a two tailed paired t-test (** = p < 0.01).

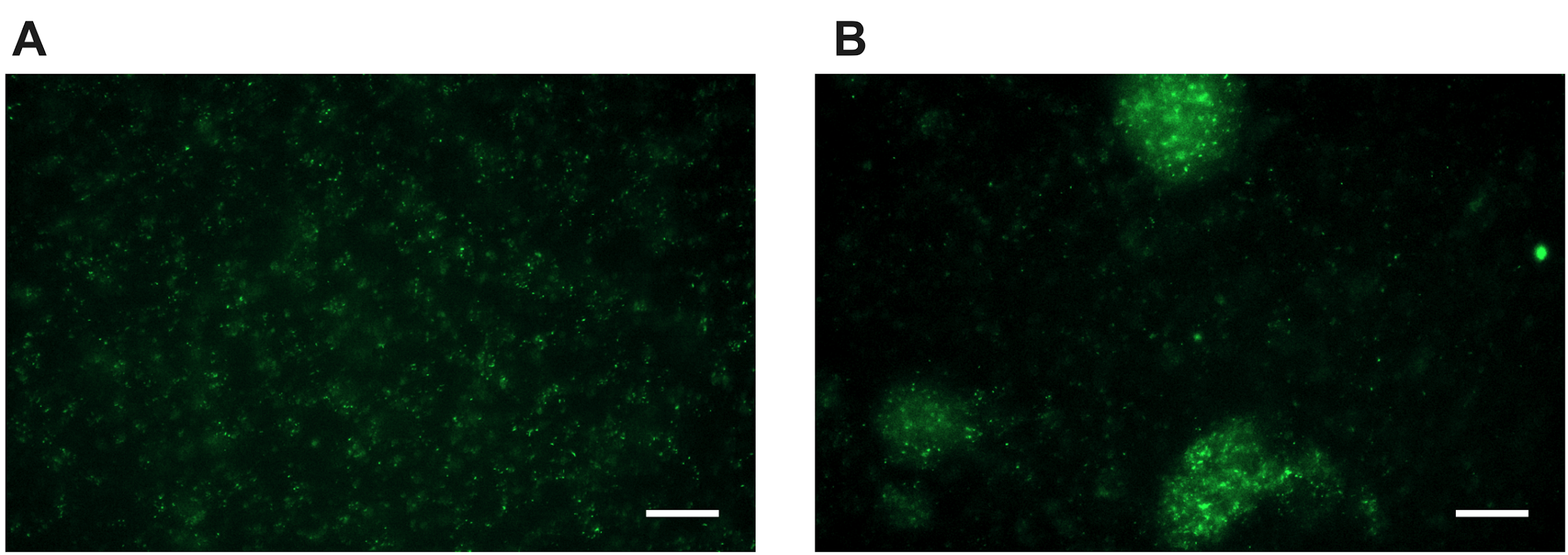

**Figure S2.**  Images of GFP-expressing PAO1 bacteria cultured in the absence of mucus for 1 hour at 37°C in A) buffer alone (control) or B) in a suspension of 1.5 mg/ml sNETs. Scale bar, 25 µm.
